# Contributions of individual satellite cells to muscle regeneration assessed using a confetti mouse model

**DOI:** 10.1101/2023.08.25.554761

**Authors:** Hans Heemskerk, N. Suhas Jagannathan, N. Jannah M. Nasir, Binh P. Nguyen, Keshmarathy Sacadevan, Paul T. Matsudaira, Peter T.C. So, Lisa Tucker-Kellogg

## Abstract

Insufficient regeneration is implicated in muscle pathologies, but much remains unknown about the regenerative output of individual muscle stem cells, called satellite cells (SCs). Prior work showed that individual SCs contribute to regeneration of more than one muscle fiber (“fiber-crossing”) after full-muscle damage. We investigated whether fiber-crossing also occurred in peripheral regions of a localized muscle injury. To assess fiber-crossing with a minimum number of mice, we used lineage tracing with confetti fluorescence, and developed a novel stochastic modeling method to interpret the ambiguity of multi-color fluorescent lineage tags. Microscopy of the regenerated muscle showed that adjacent fibers often expressed the same-colored tags. Computational analysis concluded that the observed color patches would be extremely unlikely to occur by chance unless SCs contributed myonuclei to multiple adjacent fibers (26-33% of SCs contributing to at most 1-2 additional fibers). Interestingly, these results were similar across the different regions studied, suggesting that severe destruction is not required for fiber-crossing. Our method to assess fiber-crossing may be useful for future study of gene and cell therapies that use fiber-crossing to aid muscle regeneration.

## Introduction

Satellite cells (SCs) are the resident stem cells in skeletal muscle (Relaix & Zammit, 2012), and each SC resides inside the basal lamina of its associated myofiber. Under normal circumstances most SCs are quiescent, but upon injury they are activated to produce a large pool of myoblasts, initiating the process of regeneration. Each SC is associated with (is a “satellite” of) a fiber, and SC-derived myoblasts often contribute to regeneration of that immediate fiber. However, fiber-crossing can occur when myoblasts originating from one fiber’s SC contribute to regeneration of other myofibers. Fiber-crossing is important because it could allow the therapeutic benefits of cell therapy, gene therapy, or SC-targeted drugs to be provided not only to the recipient myofiber but also to neighboring fibers (Goldstein *et al*, 2019). Such phenomena may determine the success of therapeutic interventions for sarcopenia, muscular dystrophies, etc.

Fiber-crossing has been documented to occur with revertant fibers in Duchenne muscular dystrophy (DMD). Revertant fibers carry a (partly) functional copy of the dystrophin gene, which makes them less susceptible to damage upon contraction. Although revertant fibers start out as singular fibers in the muscle, the revertant territory becomes larger with age, and patches of adjacent revertant fibers are formed (Lu *et al*, 2000; Yokota *et al*, 2006), suggesting that healthy myonuclei cross the (perhaps damaged) basal lamina and contribute to repair of neighboring fibers. This patchiness is seen in both mice and in human patients (Lu *et al*, 2000; Yokota *et al*, 2006; Fanin *et al*, 1995).

A seminal discovery concerning fiber-crossing was obtained by lineage tracing experiments in young and aged mice, which showed that resident SCs can indeed contribute to regenerating multiple fibers (Tierney *et al*, 2018). This study caused homogenously severe injury to the tibialis anterior and extensor digitorum longus, where the muscle tissues were “stabbed” 10 times and injected with barium chloride (BaCl_2_) in several locations to ensure its even distribution (Tierney *et al*, 2018). After severe injury, damaged fibers get removed by the immune system, resulting in a need for new fibers to be regenerated *de novo*. Furthermore, a significant number of SCs are killed after a homogeneously severe BaCl_2_ injury (Hardy *et al*, 2016), which alters both the availability of SCs and the burden on remaining SCs to contribute to regeneration (Kottlors & Kirschner, 2010). In contrast, many human muscle pathologies (e.g., DMD) arise from milder injuries that occur repeatedly. Mild injuries have a less extensive inflammatory phase, less SC death per injury, and can be repaired by fusing with myoblasts or with a newly formed myotube. Because different injuries create different challenges to muscle regeneration, we used a different injury, cardiotoxin injection, to assess fiber-crossing and to study the regenerative capabilities of SCs.

To track the contributions of individual SCs during muscle regeneration, we used a mouse model in which the satellite cell lineage is traced by inducible expression of four fluorescent “confetti” colors (Tierney *et al*, 2018; Snippert *et al*, 2010). We then use this mouse model to study 16-day regeneration after a single-dose injection of cardiotoxin in adult mice, similar to our earlier study looking at acute cardiotoxin injury in the panniculus carnosus muscle (Nasir *et al*, 2023). The single injection of cardiotoxin generates localized injury, giving us the opportunity to study central, peripheral, and intermediate regions of the injury. After regeneration, our microscopy of muscle fiber cross-sections showed that each region of injury displayed “iso-chromatic patches”, meaning adjacent myofibers with the same color. Iso-chromatic patches create a fundamental uncertainty of analysis, because they can arise if multiple fibers are regenerated from the same SC (therefore bearing the same color), but can also arise if regenerated from multiple different SCs that happened to have the same fluorescent color by chance.

Using the confetti mouse creates an analysis need, because 4 colors is not enough for every SC to have a different color. We know that if adjacent fibers have different colors, then they must have been generated from different SCs. However, if adjacent fibers have the same color, then we cannot be sure whether they were generated by the same SC (as opposed to being generated by different SCs having the same color by chance). We developed statistical simulations to estimate the likelihood that the observed color patches arose by chance, meaning that the color(s) of each fiber would be generated independently of neighboring fibers. The statistics show that individual SCs must have contributed myoblasts to multiple fibers, even in the peripheral (least injured) region of the muscle.

## Results

### Baseline turnover

The transgenic mouse model (Tierney *et al*, 2018; Nasir *et al*, 2023) was validated (Supplemental Text 1, Fig. S1) and we measured the background rate of muscle repair by counting the number of fluorescent fibers at 25 days after the last tamoxifen injection. In contrast to prior studies showing high rates of myonuclear turnover in the absence of experimentally-induced injury (Keefe *et al*, 2015; Pawlikowski *et al*, 2015), we detected no fluorescent fibers (negative data not shown). However, with a longer induction time of 1.5 years, fluorescence was very extensive (Fig. S2). One possible explanation is that in adult mice, fibers repaired by a single myonucleus might contribute undetectable levels of fluorescence when viewed in the overall fiber (due to dilution), but that such fluorescence would become detectable when multiple repair events accumulate.

### Expression of confetti colors in injured muscle

To study muscle regeneration after injury, we examined cross sections of tibialis anterior muscles isolated sixteen days after a single cardiotoxin injection (Fig. 1A). We found large regions in which most fibers were fluorescent (Fig 1B), and sometimes, the neighboring extensor digitorum longus muscle also contained fluorescent muscle fibers (data not shown). We found muscle fibers expressing zero, one, two, three or four colors of fluorescence (Fig. 1C-G). Fusion into myotubes had finished, and fusion/maturation of myotubes into myofibers was finished in large parts of the muscles. Further, we found the myonuclei expressing GFP^nuc^ to be located in both the central and peripheral locations of the fiber (Fig. S3A and B, arrowheads). In addition to fluorescent muscle fibers, we also detected some single cells with fluorescence (Fig. S3C and D, arrows), which had size and shape resembling the morphology of satellite cells.

**Figure 1:**
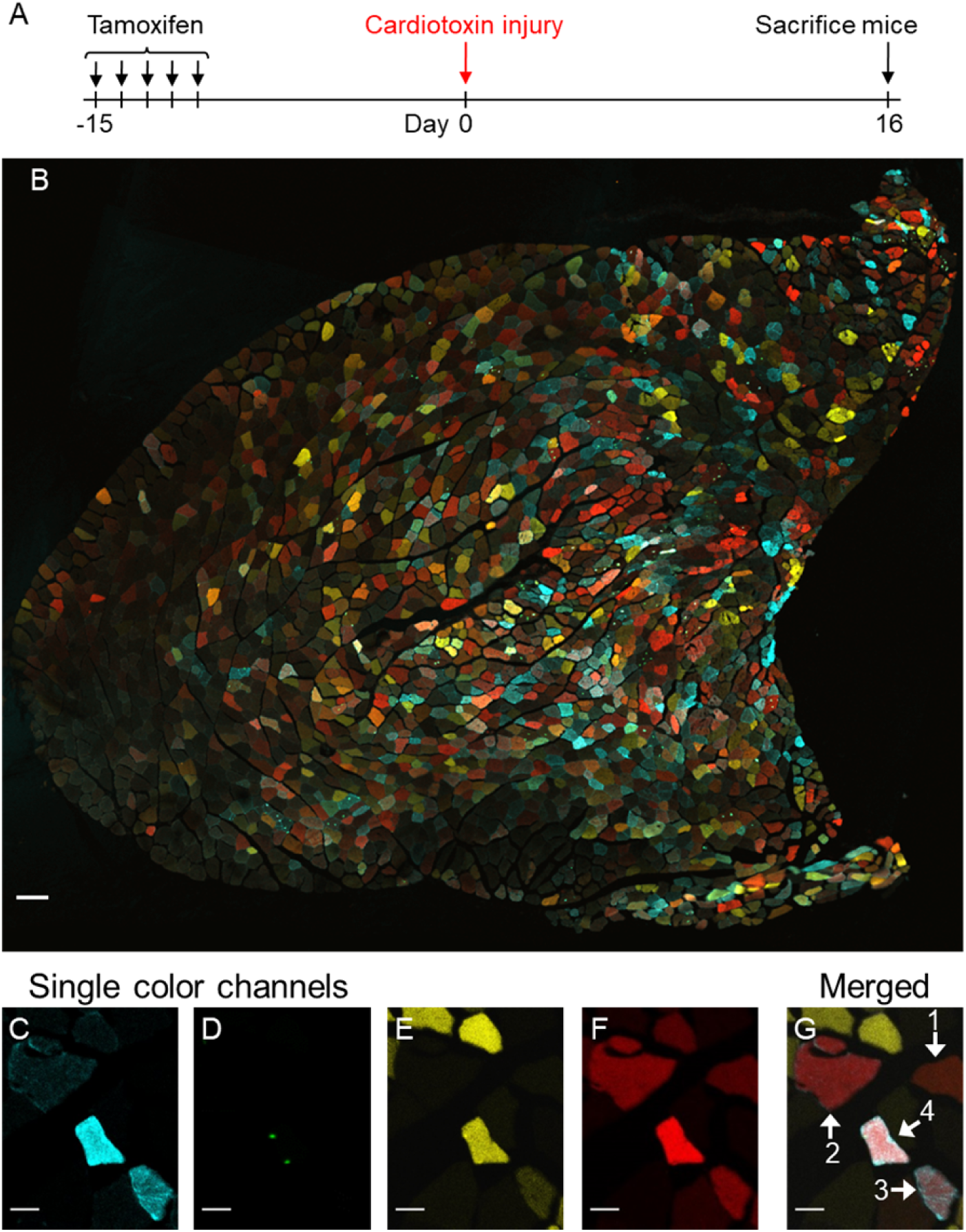
Four confetti colors are brightly expressed in regenerated muscle 16 days after cardiotoxin injury. (A) Experimental schedule, with confetti recombination induced prior to a single injection of cardiotoxin. (B) A complete cross section of tibialis anterior, at 16 days after injury, merged from 4 channels of fluorescence. (C-F) Single-color channels and (G) 4-channel merge of the same field of view, indicating that some myofibers are single-colored and some multi-colored. Numbered arrows indicate fibers expressing 1, 2, 3, or 4 colors. Scale bars are 100µm (B) or 20µm (C-G).

### Muscle areas at different stages of the regeneration process

The cardiotoxin injection created a focal region of injury with the center of the tibialis anterior having a greater extent of injury (Fig. S4). Histology showed a gradient of both tissue disruption (Fig. S5A-C) and immune infiltration (Fig. S5D), with least amount in the periphery and increasing levels in intermediate and central regions of the injured tissue. Fiber diameters were spatially organized (Fig. 2A-F; Fig. S5E-F) with enlarged fibers located at the periphery of the injury and the smallest fluorescent fibers near the center. This is consistent with prior work (Hardy *et al*, 2016) showing obvious qualitative differences between peripheral regions with advanced stages of regeneration, versus central regions with debris, immune cell infiltrate, and less advanced regeneration i.e. small and incompletely-fused myofibers (Fig. S5E-F). Given the marked differences in fiber diameters in each region (Fig. 2G-I), we used the average fiber diameter to categorize each microscopy image as belonging to central, intermediate, or peripheral regions. The central region was defined as images with average fiber diameter <80% of the contralateral, uninjured muscle; the intermediate region was defined as images with average fiber diameter between 80-100% of the contralateral; and the peripheral region was defined as images with average fiber diameter greater than the contralateral. The mean fiber sizes were 24.2 ± 10.8 µm for central, 31.6 ± 11.7 µm for intermediate, and 38.8 ± 12.7 µm for peripheral (Fig. 2G-I). The morphology and fiber diameter of the central and intermediate areas were comparable to previously published images for this duration after cardiotoxin injury (Hardy *et al*, 2016; Mahdy *et al*, 2015). In all three regions and particularly in the central region, we observed “twin fibers,” meaning pairs of fibers having close proximity, extensive contact, and the same color. Most twin fibers had a *convex hull* that was rounded in shape, suggesting that if regeneration had continued, they could have fused together.

**Figure 2:**
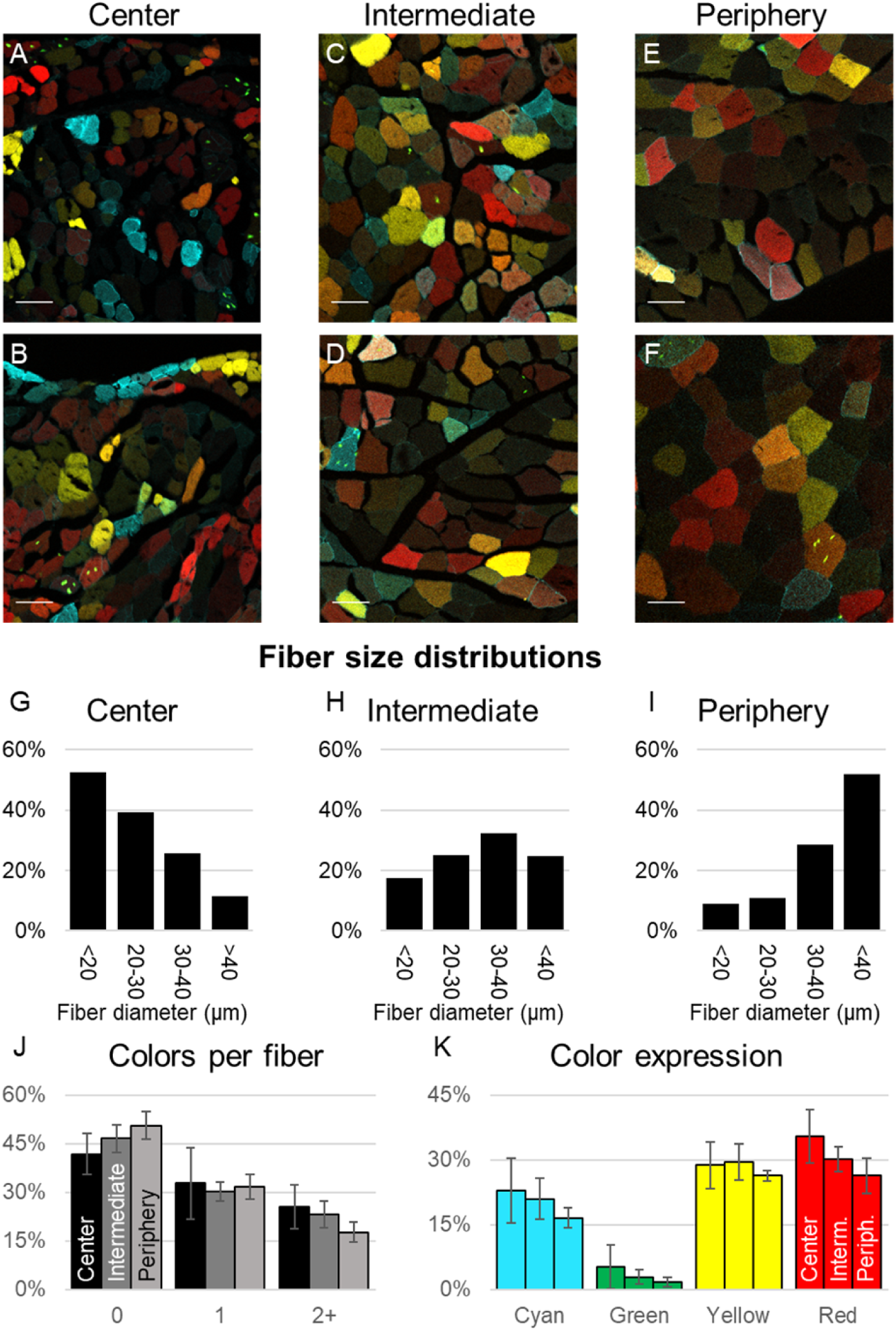
Subdivision of injured area into central, intermediate and peripheral regions. (A-F) Representative images from the central (A and B), intermediate (C and D) and peripheral regions (E and F). (G-I) Histograms of the fiber size distributions in the three regions. (J) Percentage of fibers (by region) expressing no fluorescence, 1 color and more than 1 color. (K) Percentage of fibers (by region) expressing each of the four confetti colors. (N=4, scale bars are 50µm).

We next analyzed the number of fibers expressing each of the 4 colors (Fig. 2J-K), bearing in mind the low abundance of CFP^mem^ and GFP^nuc^ cells in uninjured tissue, and bearing in mind prior reports that GFP^nuc^ expression may be suppressed. In our regenerated regions, there was strong fluorescence of many colors, but the number of fibers expressing GFP^nuc^ (3%) was approximately 10 times lower than the number of fibers expressing CFP^mem^, YFP^cyt^ and RFP^cyt^ (18-32%).

### Analysis of iso-chromatic patches

In all three regions, we frequently observed clusters of adjacent fibers having the same color (“iso-chromatic patches,” Fig. 3A-D). Iso-chromatic patches were most striking for GFP^nuc^: even in micrographs with extremely few fibers expressing GFP^nuc^, many of the GFP^nuc^-positive fibers were adjacent to each other (Fig. 3B).

**Figure 3:**
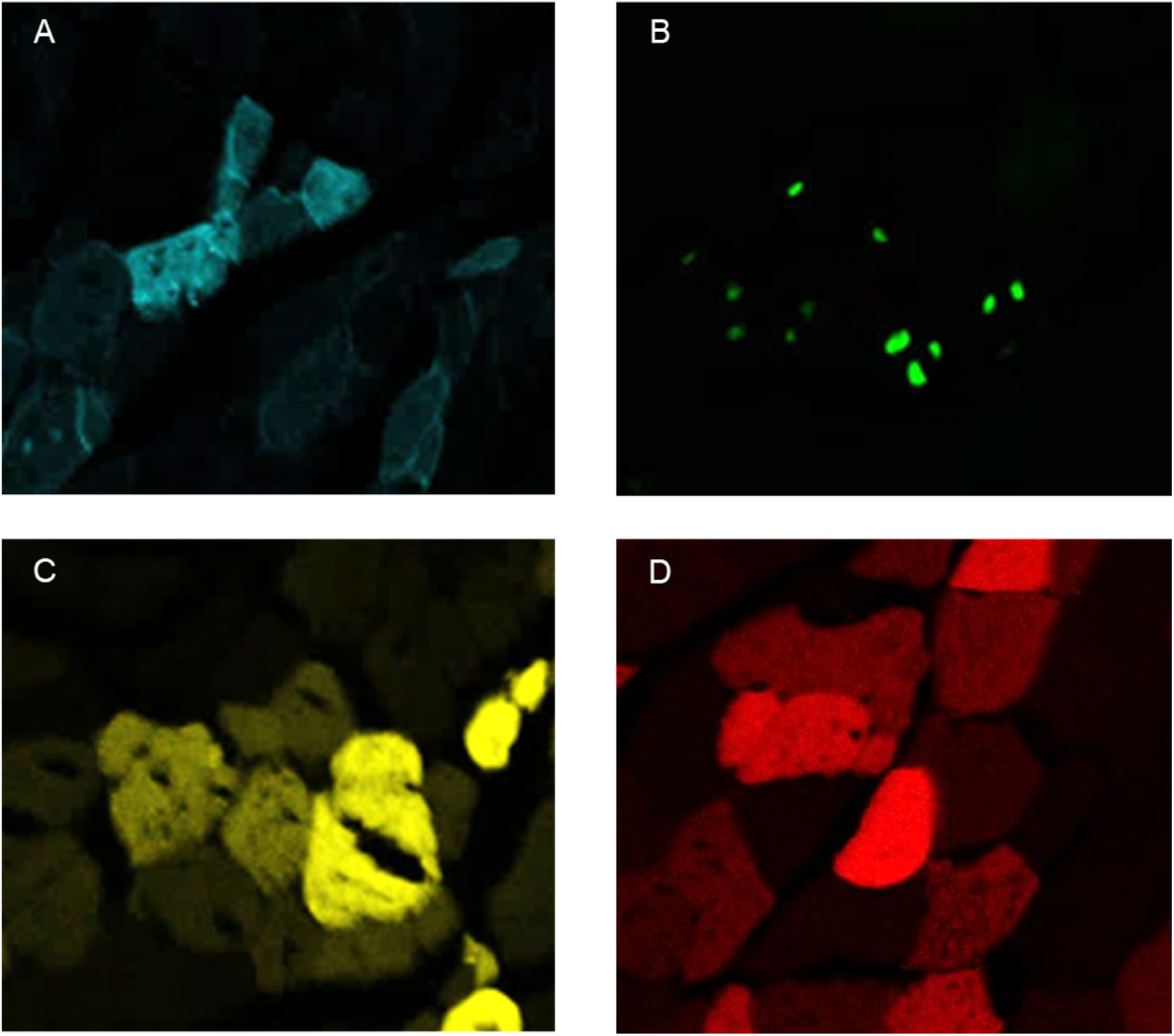
Patches of neighboring fibers with the same color. (A-D) Examples of iso-chromatic patches, for each of the four colors.

The presence of iso-chromatic patches suggests that fiber-crossing may have occurred. However, iso-chromatic patches are non-intuitive to interpret, because whenever a patch contains two or more fibers, we do not know how many different SCs created it. Two singleton fibers (fibers with no neighbors sharing the same color) or small patches generated by different SCs (that happen to have the same color by chance) could occur beside each at random, resulting in the observation of a single larger patch instead of two smaller ones. Although such questions can never be answered conclusively for a particular patch, the bulk fraction can be determined with high confidence for a statistical population. In other words, we can determine whether fiber-crossing has occurred by asking the following question: If it were true that each SC contributed to only one fiber, how likely (or unlikely) would it be, to obtain the patch size distributions (frequency of patches of different sizes) observed in our micrographs? If the analysis shows it would be extremely unlikely for the observed patches to possibly arise from SCs that can only contribute to their own fiber, then we will be able to conclude that fiber-crossing occurred.

To answer this question, we implemented the following stochastic modeling algorithm, that we hereafter call the Monte Carlo redistribution algorithm (MCRA). First, the algorithm detected and quantified the iso-chromatic patches in each of the original micrographs as follows. We obtained segmented fibers for each cross-sectional image, using our previously developed computational pipeline (Supplemental Methods figure 1F) (Nguyen *et al*, 2016). We then employed a semi-automated procedure that used fiber centroids and distances to identify *adjacent* fibers (i.e., neighbor-neighbor relationships) (Supplemental Methods figure 1H). Using this network of adjacency relationships, we extracted subnetworks corresponding to each confetti color (containing only those fibers that displayed the color). From the subnetwork for each color, we then delineated all distinct patches of that color (in formal terms, we identified the connected components of the adjacency graph for each color). Finally, from the identified patches, we obtained patch size distributions in these original micrographs.

To quantify our expectations, we computed the statistical distribution of iso-chromatic patch sizes (the “patchiness”) that would be expected if the null hypothesis were true. This is the histogram of patch sizes if one assumes that the colors of each fiber are independent of the colors of neighboring fibers. Note that this distribution is non-trivial to compute because the likelihood of duplicate colors appearing beside each other by chance is dependent on the observed fiber geometry as well as the color frequencies. For example, if a fiber has more neighbors, then the expected value of its patch size would be larger; and if a fiber has an infrequently expressed color such as green, then the expected value would be smaller. The quantified expectations were computed as follows. First, to avoid counting immature myotubes as separate fibers, we merged “twin fibers” in the intermediate and peripheral region into one fiber. In the central region, “twin fibers” were too numerous to merge and their morphologies were too ambiguous. Therefore, to be conservative and to avoid the risk of over-estimating the occurrence of fiber-crossing, we omitted the central region from further analysis. Next, we counted how many fibers in our images were positive for each of the 4 colors. Finally, we created a series of 1000 randomly-scrambled color assignments for our images, according to the null hypothesis. The MCRA scrambling process creates pseudo-images that maintain the same fiber geometry and adjacency, but randomly shuffle which colors occur where. The scrambled color assignments maintained the same total color frequency as in the original images (see Methods). We chose to redistribute the observed colors, rather than generating color assignments *de novo*, because a redistribution approach does not require any additional normalization to account for non-ideal expression levels, nor correction for unequal expression frequencies. In other words, shuffling creates a distribution that (by definition) contains the same skews as the experimental observation. During the MCRA scrambling process, fibers are not restricted to having just one color, consistent with the observation of multi-color fibers in the original cross-sectional microscopy. Multiple colors can result from fluorescent molecules mixing along the fiber axis from out of the image plane.

For each MCRA-generated pseudo-image and each color, we computed the iso-chromatic patch sizes. Thus, randomization provided expected distributions for the patch sizes that would occur if the null hypothesis (no fiber-crossing) were true. We then compared these expected distributions vs. observed frequencies for the two extremes of patch sizes: singletons (patches of size 1), and mega-patches (patches at or greater than the 85th percentile for patch-size). Colored histograms in Fig. 4 show the expected frequency distributions of singletons and mega-patches in the peripheral region (for each color), when assuming that no fiber crossing occurs. The black vertical line in each panel shows the observed fraction of fibers in iso-chromatic patches of a given size (singletons or mega-patches). The black vertical lines fall far from the expected distributions in a consistent manner: mega-patches are far more common than expected, and the singleton fibers are far less common than expected under the null hypothesis. This indicates that our null hypothesis (no fiber-crossing) is very unlikely in the peripheral region. The peripheral region is our chief interest for normal muscle physiology because SC death is expected to be low in this region and because many fibers are repaired rather than regenerated de novo. As expected, the intermediate region with more severe injury also refutes the null hypothesis and exhibits fiber-crossings (Fig. S6).

**Figure 4:**
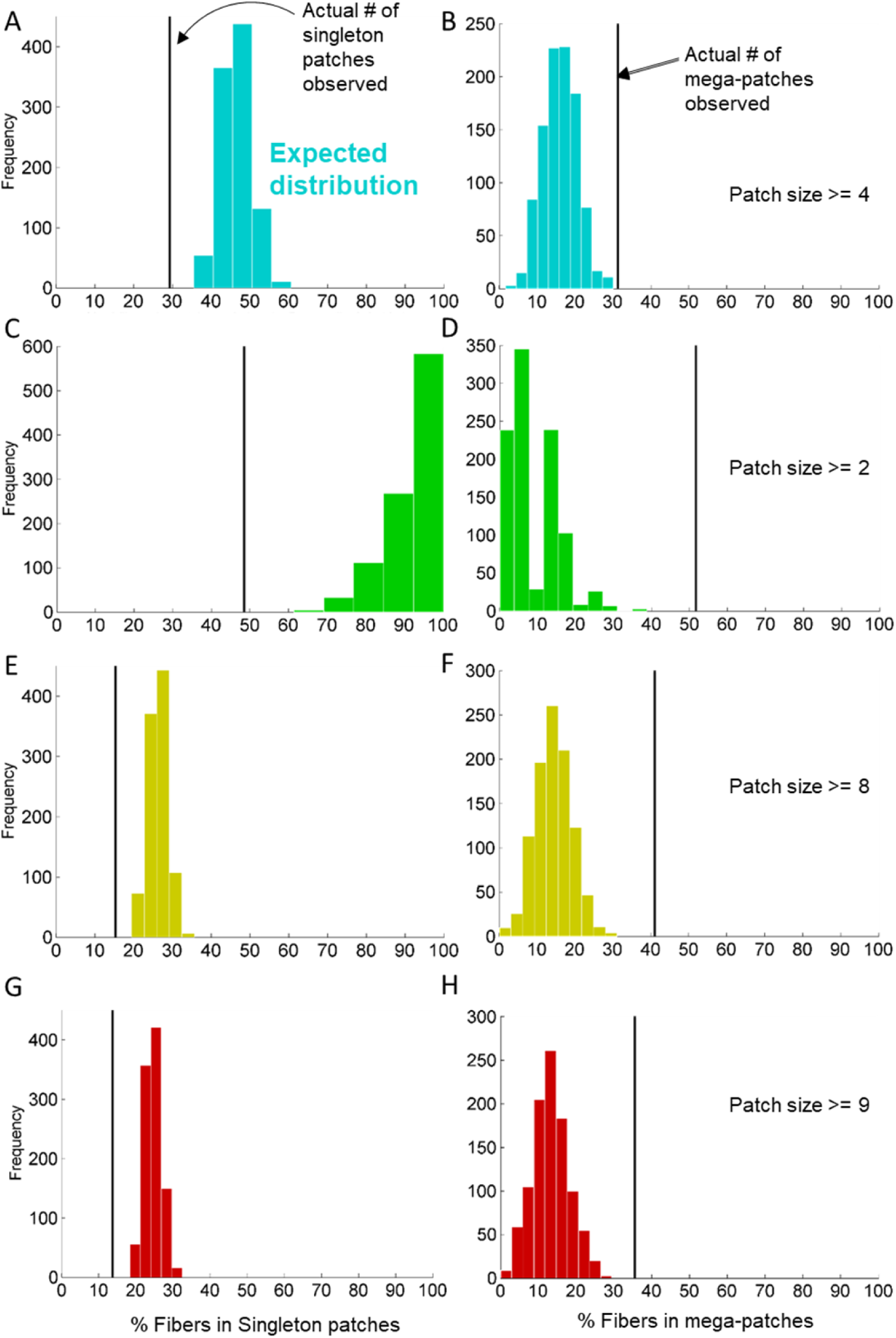
Iso-chromatic patch sizes in the peripheral region of the injury. (A-F) Comparison of patch data from experiments (vertical line) with expected patch size distributions (colored histograms), which were calculated by assuming independence between neighboring fibers and no SC fiber-crossing. Fibers not part of a patch (singletons) were far less prevalent in the experimental data (A, C, E, G) than in the expected distributions. Conversely, the number of fibers in the largest patches (mega-patches) were far more prevalent in the experimental data (B, D, F, H) than in the expected distributions. The threshold for defining mega-patches was determined for each color as explained in the Monte Carlo confetti redistribution algorithm section of Methods.

To confirm that the divergence between observation and expectation was not dependent on the thresholds we used for categorizing fibers, we repeated the entire analysis using more conservative thresholds for delineating the region of mildest injury (Fig. S7). In all versions, comparing the observations against the expected distributions disproved the null hypothesis. Therefore, the fluorescent myoblasts that contributed to the observed regeneration were non-independent across adjacent fibers. We conclude that at least some of the individual Pax7+ cells from the time of tamoxifen induction must have contributed to regeneration of multiple fibers.

### Estimate of the extent of migration across fibers

From earlier results, we inferred that SCs do cross the territorial boundaries typically demarcated by basal lamina – in other words, passing through ruptures in the basal lamina or crossing the basal lamina. We next sought to quantify the extent of SC migration required to recapitulate observed patch size distributions. Specifically, we asked what fraction of labeled-SCs would be required to migrate, and how many additional fibers these SCs contribute to, on average. To answer this question, we repeated our Monte-Carlo confetti redistribution algorithm (MCRA), but this time with the following assumption: the number of additional fibers each SC contributes to (in addition to their home fiber) would follow a Poisson distribution with mean λ. We performed simulations using different values of λ and estimated the value of λ that best fits the experimental observations for each color and region (Fig. S8 and S9, and Table S1). The overall best-fit estimates of λ (across all colors) were found to be 0.4 and 0.3 for the peripheral and intermediate regions, respectively. This corresponds to having roughly ∼33% of SCs in the periphery, and ∼26% of SCs in the intermediate region, contributing to at most 1-2 additional fibers.

## Discussion

In this study, we used confetti labeling to assess the contribution of Pax7+ cells to muscle regeneration, and we developed image analysis and computational methods to process the generated data. After cardiotoxin injury, microscopy revealed bright fluorescent colors in the regenerated muscle (compared with black or nearly black surrounding region, which we interpret as uninjured or nearly uninjured). All four colors had bright fluorescence, even cyan and green, but green was very infrequent. One explanation for the low frequency of GFP^nuc^ is that recombination of the transgene sometimes results in the absence of color instead of GFP^nuc^ expression (Tarlow *et al*, 2014). Another consideration is that GFP^nuc^ is confined to nuclei, while cytoplasmic (YFP^cyt^ and RFP^cyt^) or sarcolemmal (CFP^mem^) labels are expected to diffuse along the fiber (Chretien *et al*, 2005). Being spread out along the axis would allow those colors to be visible in more fibers when viewed as cross-sectional images.

Although the regenerated muscle showed bright fluorescence in many fibers, some fibers were dim or non-fluorescent. Non-fluorescent fibers were detected throughout the injured muscle and can be interpreted in 3 ways. The first possibility is that the fiber was regenerated from Pax7 progenitor(s) but no fluorescence was expressed, as seen previously (Tierney *et al*, 2018). This could arise from failed recombination in the SC, most likely for GFP^nuc^ (Tarlow *et al*, 2014). The second possibility, suggested by the ability of cardiotoxin to kill SCs (Hardy *et al*, 2016), is that the fibers may have regenerated from non-Pax7 progenitors. The final possibility is that non-fluorescent fibers may have survived the cardiotoxin without needing significant regeneration. Some of the non-fluorescent fibers had peripheral location of nuclei, supporting this interpretation.

When we quantified fiber diameters, we found a gradient with smaller fibers toward the center of the injury and larger fibers toward the periphery. We attribute this to different levels of cardiotoxin exposure and different degrees of tissue damage. We categorized the regions into central, intermediate, and peripheral. In the peripheral region, regeneration was nearly complete at 16 days, and also at 8 days. Furthermore, the large fiber diameter seen in the periphery at 16 days is comparable to the fiber diameters seen after milder injuries (e.g., large-strain lengthening contractions (Roche *et al*, 2008) instead of homogeneous injections of cardiotoxin or BaCl_2_). We interpret that the peripheral region received low cardiotoxin exposure, and that many of the peripheral fibers acquired fluorescence via repair rather than replacement. We speculate that this region would have less death of SCs. Beyond the peripheral region is dark (non-fluorescent) tissue, which we interpret as uninjured. In the intermediate region, located between central and peripheral, regeneration was more advanced than in the central region, and “twin fibers” were less frequent. Additional samples taken at 8 days post-cardiotoxin indicate that many fibers in the intermediate region have been killed. Levels of SC death are unknown, but prior work showed high levels of SC death (Hardy *et al*, 2016) in a model of cardiotoxin injury that might resemble our central and intermediate regions. In the central region, thin myotubes and “twin fibers” were still present at 16 days, indicating incomplete regeneration. This is consistent with the central region receiving the highest concentration of cardiotoxin and the greatest severity of injury. In subsequent discussion, we interpret the peripheral, intermediate, and central regions as moderate, severe, and extreme injuries, but the nature of the gradient requires future work to fully characterize.

The most intriguing result for confetti-SC regeneration was that the fluorescent colors sometimes occurred in clusters, which we called “iso-chromatic patches.” To ascertain whether the mixing of colors was random (i.e., whether the clustering was statistically significant), we developed image analysis tools and used stochastic modeling. With these tools we show that the observed size and frequency of iso-chromatic patches is statistically incompatible with the assumption that the colors of each fiber are independent of the colors of its neighbors. We therefore conclude that some non-independent process must have occurred between neighboring fibers. Since recombination of the confetti transgene is stochastic, we interpret non-independence to indicate that some of the SC pool contributed myonuclei to two or more adjacent muscle fibers. Additional analysis suggested the patches could be explained if ∼26-33% of the SCs contributed to one or more additional neighboring fibers.

SCs have previously been shown to contribute myoblasts to neighboring muscle fibers, but the damage caused by severe injury could have facilitated fiber-crossing (Tierney *et al*, 2018). Here, we found that fiber-crossing occurred to a similar degree in both the peripheral and intermediate regions, despite the differences in injury severity. This suggests that fiber-crossing might be possible during fiber-repair, not just during de novo regeneration; moreover, it might not require severe inflammation. Our analysis does not disambiguate whether the myonuclei were transmitted via migration of myoblastic progenitor cells, creation of new SCs, or other cell types. Future work should track additional cell types or architectural features (e.g., membrane rupture) that contribute to fiber-crossing.

There are several challenges that can impair SC regeneration of muscle fibers. Proliferative output can be low or delayed, for instance if SCs are in a state of deeper quiescence (Rocheteau *et al*, 2012). Functionality can be reduced due to cellular alterations such as senescence (Sousa-Victor *et al*, 2014), or signals in the niche (Chakkalakal *et al*, 2012; Lacraz *et al*, 2015), or in the systemic environment (Brack *et al*, 2007). These challenges reduce the efficiency of the regeneration process and can lead to fiber loss and an increase in fibrosis. Highly proliferative SCs might rescue regeneration in these fibers, if they can supply cells through fiber-crossing. Therefore, it is important to measure and analyze fiber crossing.

Another reason to analyze fiber crossing is to assess the beneficial impact of stem cell (Negroni *et al*, 2015) and gene therapy (Duan, 2018; Lim *et al*, 2018; Chemello *et al*, 2020; Min *et al*, 2019; Zhang *et al*, 2018). The eventual impact of gene therapy and stem cell therapy could be greater than the number of SC affected, if fiber-crossing permits delivery of a therapeutic intervention from a resident SC to neighboring fibers. In particular, CRISPR/Cas9 based gene therapy has shown promise toward mitigating muscle disorders such as Duchenne muscular dystrophy (DMD) by increasing dystrophin positive muscle fibers through the engraftment of gene-edited Pax7+ stem cells both in uninjured muscle (Tabebordbar *et al*, 2016) or after serial injury (Kwon *et al*, 2020). In mice models of DMD (mdx mice), transplantation of small amounts of muscle satellite cells in the tibialis anterior muscle (20–200 MuSCs/mg muscle) showed a marked increase in muscle function (Cosgrove *et al*, 2014; Judson & Rossi, 2020). In such cases, a combination of therapeutic gene editing, confetti fluorescence, and stochastic modeling (MCRA) could be used to quantify multiple parameters, such as the starting pool of SCs required for transplantation, extent of fiber crossing by SCs in DMD, and the persistence of SCs and their ability to provide a self-renewing source of corrected muscle fibers over time. Thus, our approach provides a novel method to assess fiber-crossing of genetic material in myofibers, similar to (Tierney *et al*, 2018), without requiring a large number of mice.

Research on revertant fibers in DMD mouse models suggests there might indeed be a beneficial effect of fiber-crossing. When different strains of mice are grown with the *mdx* model of disease, the strain with less fiber-crossing (i.e., smaller patches of revertant fibers) have more severe disease (Fukada *et al*, 2010; Rodrigues *et al*, 2016). The same strain of mice have greater muscle loss during aging (Lionikas *et al*, 2006). However, it is not known whether lack of fiber-crossing is a cause or a consequence of these muscle pathologies. The difference between mouse strains also indicates that the level of fiber-crossing could differ greatly between people. Future research should determine whether fiber-crossing is an active process, as has been suggested by experiments on isolated muscle fibers (Collins-Hooper *et al*, 2012), or if it is a by-product of inflammation and basal lamina disruption. Additionally, future studies should determine whether fiber-crossing contributes to undesirable outcomes such as progression of rhabdomyosarcoma, or formation of non-functional muscle fragments in the interstitium (Pisconti *et al*, 2016). Finally, repeated severe injuries have been shown to lead to clonality in an increasingly large area (Tierney *et al*, 2018). Cells might have divided several times before crossing over to a neighboring fiber, and therefore, newly created SC in the destination might have shorter telomeres than the original SC. Telomere shortening leads to senescence, and interestingly, before SCs go into senescence they display a lower level of fiber-crossing (Sacco *et al*, 2010). Therefore, it would be interesting to determine whether, in cases of repeated injuries, any beneficial effect of fiber-crossing would be limited to a certain number of degeneration/regeneration cycles.

Our study has some caveats for both the experimental and computational approaches. By using destructive analysis of samples, we rely on specific timepoints rather than using non-invasive longitudinal imaging to track dynamic trends prior to sacrifice. Our study ended at 16 days (in keeping with prior studies of post-cardiotoxin myogenesis (Goetsch *et al*, 2003), but it is possible that analysis of a later timepoint could reveal greater levels of lateral fiber crossing that are not yet visible at 16 days. We also do not attempt to infer the number of rips or ruptures in the basal lamina, and this is likely to be important for the migration of progenitor cells into neighboring fibers. The cardiotoxin injury used in this study causes some degeneration and de novo fiber formation, in a gradient of severity. For a more definitive study of mild injury, a different type of injury could be used, that does not cause de novo fiber formation. Another limitation of our analysis concerns the image analysis, because the fiber adjacency networks are first extracted by the algorithm and then refined manually. The refinement process is under the control of a human operator and if the operator behaves inconsistently, could create minor differences in fiber neighbor relationships. On the computational side, one weakness of our MCRA algorithm is that it cannot differentiate the background of uninjured fibers (which do not express any fluorescent color) from the injured-but-uncolored fibers in the injured region that lack fluorescence due to failed recombination in the respective SCs. Although this skew is accounted for by the MCRA (and therefore does not affect the relative comparison between observed images and shuffled pseudo-images), it may nonetheless affect the estimation of absolute parameters such as λ, the number of additional fibers each SC may contribute to. The estimated value of λ is also subject to our assumption that λ follows a Poisson distribution.

In conclusion, we show here that fiber-crossing to neighboring muscle fibers occurs in peripheral regions, not just in severely injured regions of injury. Fiber-crossing could ensure efficient regeneration when an SC on a neighboring fiber does not contribute sufficiently to regeneration. It could also be important for cell-based therapies and gene therapies to spread therapeutic effects over a larger area.

## Resource Availability

Further information and requests for resources and reagents should be directed to and will be fulfilled by the Lead Contact, Lisa Tucker-Kellogg. The mice used in this study were purchased from Jackson Laboratories and Cyagen Biosciences. The image analysis software used in this study is available in the journal article published by Nguyen et. al (2016) (Nguyen *et al*, 2016). The software for computing expected cluster sizes is available at https://github.com/nsuhasj/MCRA.

## Methods

### Mice

Animal experiments were approved by the institutional animal care and use committee of the National University of Singapore. To conditionally label the Pax7 expressing cells, we crossed the Pax7-Cre-ER^T2^ with the R26R-Confetti mouse, similar to our earlier study (Nasir *et al*, 2023). The Cre-ER^T2^ fusion protein is located downstream of the Pax7 stop codon and will only be expressed in cells expressing Pax7. Additionally, it is inducible by tamoxifen and therefore inactive until tamoxifen is administered to the mice. Once the Cre-ER^T2^ is activated, it will recombine the confetti construct, resulting in the random expression of one of the fluorescent proteins in the construct. The fluorescent proteins are mCerulean (CFP^mem^), hrGFP II (GFP^nuc^), mYFP (YFP^cyt^) and tdimer2(12) (RFP^cyt^). CFP^mem^ will be localized tethered to the sarcolemma, GFP^nuc^ will be localized in the nucleus and YFP^cyt^ and RFP^cyt^ will be located in the cytoplasm.

### Muscle injury

Six-month-old mice were used for the experiments, both male and female (n = 4), and heterozygous for both Pax7-Cre-ER^T2^ and R26R-Confetti alleles. The mice were intraperitoneally injected with tamoxifen on five consecutive days (100 µg/g, Sigma, Singapore) to induce recombination. To ensure the confetti colors are only present in cells expressing Pax7, in case later events trigger additional cells to express Pax7, there was a 10 day clearance between last tamoxifen injection and the muscle injury. To induce damage to the muscle fibers, the tibialis anterior muscle of the right leg was injected with 50 µl cardiotoxin (10 μmol, Sigma, Singapore). The contralateral muscle was used as uninjured control.

### Identification of iso-chromatic patches

We developed a semi-automated process (Supplemental methods) to identify the adjacent neighbors of each fiber. The procedure used the centroids and diameters of the fibers to compile a list of candidate neighbors for each fiber. Then a streamlined graphical user interface allowed a human user to manually approve or correct the candidate neighbors. The human user also manually merged fibers that were deemed to be “twin fibers” (immature myotubes with convex outer edge, believed to be within the same basal lamina and/or in the process of fusing into a single myofiber). The network of all adjacency relationships (i.e., the union of all sets of adjacent-neighbor relationships) was represented as an adjacency matrix ***Adj***.

For each color (Cyan, Red, Yellow, Green), we define a sub-network of ***Adj*** that only contains fibers that exhibit the given color. For example, the yellow sub-network of ***Adj*** would contain all fibers that express yellow (even if they express other colors as well), and all adjacency relationships between pairs of fibers, where both fibers express yellow. We then define *iso-chromatic patches* to be the *connected components* of the color-specific sub-network. For example, the yellow iso-chromatic patches would be the groups of yellow-expressing fibers that are adjacency-connected. A yellow fiber with entirely non-yellow neighbors would be an iso-chromatic patch of size 1. Patch membership uses adjacency as a *transitive relation*, meaning that an iso-chromatic patch could be a snaking chain of connections rather than a *clique*. Note that iso-chromatic patches are defined to have maximal size, meaning they are not adjacent to any same-color fibers that are not in the set.

The computation of iso-chromatic patches was repeated and pooled for all micrographs corresponding to a region (intermediate/periphery), for 4 mice. From the identified patches, we obtained the fraction of all fibers of a particular color (e.g., yellow) that were members of a patch of size *p* (1 ≤ *p* ≤ *p_max_*) where *p_max_* represents the size of the largest observed patch in the pooled set of images for each region. Central region micrographs (severe injury) were omitted from the analysis because iso-chromatic patches would be skewed by the presence of “twin fibers”. Such myotubes are very numerous in the central region, creating ambiguity about the fusion or twinning relationships, so that “twin fibers” could not be manually annotated without introducing human bias. In contrast, the “twin fibers” of other regions were infrequent and shared a more obvious *convex hull*.

### Monte Carlo confetti redistribution algorithm (iso-chromatic patch sizes without fiber-crossing)

To obtain an *expected distribution* for the iso-chromatic patch sizes, we generated 1000 pseudo-images subject to the assumption that fiber colors are statistically independent of neighbor fiber colors (*i.e*., subject to the null hypothesis that fiber-crossing does not occur). The pseudo-images were generated from the observed images by preserving the adjacency information (***Adj***) and the total color counts (*n_C_*), while randomly scrambling the color locations, according to the following algorithm: we took an uncolored image and for each color *C* (cyan, red, yellow, green), we repeatedly picked a fiber at random and assigned it the color *C*, until the total number of distinct *C*-colored fibers equaled *n_C_*. Finally, the iso-chromatic patches were identified and counted, across 1000 random pseudo-images, to provide the expected fraction of fibers in iso-chromatic patches of size *p* (for all colors *C*, and for all patch sizes *p*). For comparison between observed values and expected distributions, we defined a *mega-patch* of each color *C,* to be any patch of size > *p_L_(C)*, such that in the observed images, patches of size > *p_L_(C)* contain ∼15% of *n_C_* (all fibers of color *C)*.

### Monte Carlo confetti redistribution algorithm (iso-chromatic patch sizes with fiber-crossing)

We first assume that each SC that contributes to regeneration is in immediate contact with one myofiber, to which it can contribute by default. Simulation then focuses on the amount of SC contribution to fibers that are adjacent-neighbors to its immediate fiber (because we assume there is no contribution to distant, non-neighboring fibers). We next assumed that the number of neighboring fibers to which an SC contributes would follow a Poisson distribution with mean λ (λ ≥ 0). Hence, the total number of fibers to which each SC contributes is 1 + Poisson(λ).

We implemented the following Monte Carlo simulations to consider possible values of λ and find which value provides the best fit the with the observed patch sizes. For each λ value considered, 1000 pseudo-images were generated as follows. For each pseudo-image, the adjacency network ***Adj***, and the total color counts (*nC*) were preserved from the observed image, while original color information for all fibers was “forgotten”. Each color *C* was iteratively assigned to a randomly chosen set of ***X*** fibers (X ∼1+Poisson (λ)), such that the set was connected, meaning all fibers in the set were neighbors of at least one other fiber in the set. This was repeated until the total number of distinct fibers displaying color *C* (*n_C_*) was the same as that in the original dataset. Finally, the iso-chromatic patches were identified and counted, across 1000 random pseudo-images, to provide the expected fraction of fibers in iso-chromatic patches of size *p* (for all colors *C*, and for all patch sizes *p*).

We searched over possible values of λ in increments of 0.05. The best-fit value λ*_best_* was then taken to be that value which minimized the sum-of-squared Z-scores computed as,

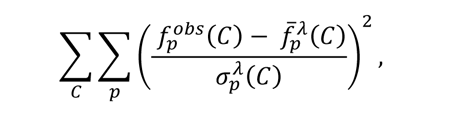

where *f*_p_^obs^(*C*) is the observed fraction of *C*-colored fibers in patch size *p*, and *f*_p_^*λ̅*^(*C*) and *σ*_*p*_^*λ*^(*C*) are the mean and standard deviations obtained using the 1000 pseudo-images.

## Acknowledgments

We sincerely thank Dr. Ralphe Bunte, board-certified veterinary pathologist, for valuable advice to develop the histopathological scoring system, Ms. Emily Dean for help in quantifying centro-nucleated myofibers, and Dr. Cheung Yin Bun for helpful advice. We also thank Dr. R. Manjunatha Kini for providing naniproin, a close homologue of cardiotoxin, although it was not used in dataset published here. Lastly, we gratefully acknowledge funding from the Singapore Ministry of Health’s National Medical Research Council under the Open Fund Individual Research Grant scheme (NMRC/OFIRG/0007/2016), and from the National Research Foundation (NRF), under its CREATE programme, Singapore-MIT Alliance for Research and Technology (SMART): Critical Analytics for Manufacturing Personalised-medicine (CAMP).

## Author Contributions

Conceptualization and Writing Original Draft, HH and LTK; Formal analysis and Software, NSJ BPN and LTK; Investigation HH, NJMN, KS; Supervision LTK PTM PTCS.

## Declaration of Interests

The authors declare that they have no conflict of interest.

